# The Evolution of Fair Offers with Low Rejection Thresholds in the Ultimatum Game

**DOI:** 10.1101/162313

**Authors:** Jeffrey C. Schank, Matt L. Miller, Paul E. Smaldino

**Affiliations:** Department of Psychology, University of California, 1 Shields Avenue, Davis, CA 95616, USA; Cognitive and Information Sciences, University of California, Merced, USA

**Keywords:** cooperation, multilevel selection, fairness, altruism, ultimatum game

## Abstract

The ultimatum game (UG) is widely used in economic and anthropological research to investigate fairness by how one player proposes to divide a resource with a second player who can reject the offer. In these contexts, fairness is understood as offers that are more generous than predicted by the subgame perfect Nash equilibrium (SPNE). A surprising and robust result of UG experiments is that proposers offer much more than the SPNE. These results have spawned many models aimed at explaining why players do not conform to the SPNE by showing how Nash equilibrium strategies can evolve far from the SPNE. However, empirical data from UG experiments indicate that players do not use Nash equilibrium strategies, but rather make generous offers while rejecting only very low offers. To better understand why people behave this way, we developed an agent-based model to investigate how generous strategies could evolve in the UG. Using agents with generic biological properties, we found that fair offers can readily evolve in structured populations even while rejection thresholds remain relatively low. We explain the evolution of fairness as a problem of the efficient conversion of resources into the production of offspring at the level of the group.

**Significance Statement:** Human generosity is widespread and far exceeds that of other social animals. Generosity is often studied experimentally with the ultimatum game, in which a proposer offers a split and a responder can either accept it or cancel the whole deal. A surprising result of ultimatum game experiments is that players are much more generous than predicted while only rejecting very low offers. This has presented a theoretical puzzle, since mathematical models have generally relied on high rejection levels—just below offer levels—to maintain generosity. Using evolutionary simulations, we explain both generous offers and the rejection of only low offers as a solution to the problem of how groups can efficiently convert resources into the production of offspring.

## Introduction

The ultimatum game (UG) is a simple bargaining game that is an extensively used paradigm for investigating fairness in humans [1-4] and other primates [5-7]. In the UG, two players, a proposer and a responder, are allotted a resource to divide. The proposer offers a portion of the resource to the responder, who can either accept or reject it. If accepted, the resource is divided as proposed. If not, both players receive nothing.

Fairness in the UG can be understood in terms of Nash equilibrium strategies. A Nash equilibrium in the UG exists when the offer portion, *p*, is equal to the rejection threshold, *q*, below which a responder rejects all offers. For the UG, there are many possible Nash equilibrium strategies depending on the divisibility of a resource. For example, a fairness strategy that evenly splits a resource can be a Nash equilibrium if responders only accept offers of at least half the resource (i.e., *p* = *q* = 0.5). However, it can be shown that there is one Nash equilibrium strategy set if both players are completely rational: the subgame perfect Nash equilibrium (SPNE) for which rational responders accept the least positive offer possible from rational proposers. Strategies in which offers are greater than predicted by SPNE are considered generous and viewed as more fair than the SPNE.

Many experiments have been conducted using the UG to investigate whether human social decision making conforms to that of the idealized rational actor [1-4]. Results within and across cultures repeatedly show that the behavior of people playing the UG does *not* conform to the SPNE. Indeed, the average offer made by proposers is often quite far from the SPNE. A meta-analysis of UG experiments [2] found that proposers offer, on average, over 40% of a resource with a modal offer of 50%. These surprising empirical results have motivated a number of theoretical models that aim to explain why people behave more fairly than predicted by the SPNE.

A recent review [8] of 36 UG models classified them into six categories: alternating role-based models [9], reputation-based models [10-12], noise-based models [13,14], spite-based models [15-17] spatial-population-structure-based models [18-20], and empathy-based models [21]. All of these models provide theoretical explanations for large departures from SPNE that are at or close to an even-split Nash equilibrium fairness strategy (i.e., *p* = *q* = 0.5). A common element that runs through all of these theoretical results is that responders reject offers that are well above offers predicted by the SPNE. In other words, the models aim to explain how fair Nash equilibrium strategies can evolve far from the SPNE. If evolution favors the emergence of Nash equilibrium strategies, even those far from SPNE, then it follows that observed mean values for offers and rejection thresholds should be nearly identical. For example, mean offers of about 40% should imply that mean rejection thresholds will also be about 40%. However, this is not what the data reveal.

The meta-analysis by Oosterbeek [2] showed that the average proposer offer was over 40% of a resource, while the average rejection threshold by responders was 16% or less, and not the 40% predicted if people are using Nash equilibrium strategies to play the UG. Henrich and colleagues [4] found very similar results in a large cross-cultural study of the UG: a mean offer of 39.6% and a mean rejection threshold of 16%. Thus, not only are offers far from SPNE but people are also not using Nash equilibrium strategies. Instead, many people appear to be using strategies consisting of generous offers paired with low rejection thresholds (i.e., rejection thresholds much lower than they are willing to offer).

People also behave more fairly than expected by assumptions of self-interested rationality in simpler a game called the *dictator game* (DG) [22,4]. The DG is like the UG except that the responder cannot reject the proposer’s offer and therefore has no leverage. The obvious solution to the DG is for proposers to offer nothing to responders, but often that is not what they do in behavioral experiments [22,4]. In a previous paper on the DG [23], we used an agent-based model to show how multilevel selection can favor fairness at the level of the group when population structure is allowed to self-organize in low population density conditions. Our results demonstrated, for a range of biologically plausible parameter values, that populations could evolve levels of fairness consistent with empirical data from the DG experiments and could do so in opposition to individual selection operating against fairness. The underlying mechanism favoring non-zero offers in the dictator game is the efficient conversion of resources into the production of offspring at the level of the group. That is, when there are constraints on the conversion of resources into the production of offspring, resources can nonetheless be efficiently used at the level of the group. This can be accomplished by sharing resources that cannot immediately be converted into offspring due to those constraints on the reproductive process (e.g., gestation of offspring). They can then convert those resources into offspring and overcome the inefficiency that results from the constraints.

The basic argument is one of multilevel selection. When groups of relatively small size compete, groups that can most efficiently allocate resources among their members will increase in number through a process similar to risk pooling [24]. That is, fair resource allocation in groups results in more offspring than in groups with less fair allocation, but differs from risk pooling when resources are scarce [23]. The boundaries between groups need not be clear or precise; only statistical separation of interactions is required (as when group members interact preferentially, but not exclusively, with other members). Successful resource allocation manifests as a decrease in the variance in resources among group members, and in the context of the UG, is facilitated by more generous offers. As we previously showed in the case of the DG [23], high rejection thresholds are not required for selection to favor generosity; offers in the DG readily evolved to empirically observed levels of 28% [22]. The added presence of rejection thresholds can, as we will see, increase offers beyond rejection thresholds while these threshold values lag far behind offers—exactly what is observed empirically.

Here we show that efficient resource conversion at the level of the group can explain the empirical results of UG games, by extending the DG model in [23] to the UG. In this model, agents possessing generically biological properties repeatedly play the UG for resources to reproduce. No other theoretical assumptions about agents are built into the model. Agents are cognitively simple. They have no complex psychological features such as memory, empathy, or spite. Rather, their behavior in the UG is determined entirely by a fixed strategy (*p, q*) of how much to offer when proposing and a minimum offer to accept when responding. Our approach was to run virtual evolutionary experiments for a variety of parameter values and to discover what kinds of strategies would evolve.

## Agent-based model

We created an agent-based model in which agents are imbued with generic properties common to most organisms that engage in social behavior and in the exchange of resources, as described in [23]. These properties include *mobility* (important for engaging other agents in space), *aggregation* (a basic condition for social behavior), *lifespan, resource accumulation* (such that they can only successfully reproduce when they have sufficient resources), *reproduction of heritable traits* (including a mechanism for introducing variation), and *parental investment* (in which resources are transferred to offspring). Another important property of the model is *population structure*, in the form of groups or clusters of agents, which emerges endogenously from spatial aggregation and local reproduction of agents [23,26]. Group selection is not programmed into the model but rather multilevel selection processes emerge as population structure self-organizes from the aggregative and reproductive behaviors of agents.

Agents play the UG with spatial neighbors for resources, use those resources to reproduce offspring with their same UG strategy, move when unable to find a game partner, and die of old age. The model was coded in Java and implemented using the MASON multi-agent simulation environment [26]. Values and descriptions of the parameters described below are listed in Table S1. Figure S1 provides a flow chart of an agent’s decisions and key events at each simulation step.

Agents are located on unique cells on a 2-dimensional discrete grid with periodic boundaries. Time proceeds in discrete steps, and each agent (in random order) executes the following actions each time step (see figure S1). The agent first attempts to maintain contact with at least one other agent in its Moore neighborhood (radius *M* = 1; that is, it checks the eight closest cells for the presence of at least one other agent). We refer to such agents as neighbors. If the agent does not have any neighbors, it searches for other agents nearby, in a slightly larger Moore neighborhood (*M* = 2; that is, the 24 closest cells). If there is at least one other agent in this search area, it chooses one at random and moves toward it; otherwise, it moves to a new cell using one step of a random zigzag walk (as described in [27]). This sort of cohesion rule is common in models of social behavior such as flocking, herding, or schooling [28]. Only one agent can occupy a given cell at a time: so if an agent attempts to move into an occupied cell, the attempt fails and the agent remains in its initial location.

Each agent *i* is defined by a strategy tuple (*p*_*i*_*, q*_*i*_), denoting its offer (0 ≤ *p*_*i*_ ≤1) when playing as proposer, and its rejection threshold (0 ≤ *q*_*i*_ ≤1) when playing as responder, respectively. An agent can only obtain resources for reproduction by playing the UG with a neighbor who has not yet played during that round of play. If such a neighbor is available, the focal agent initiates a round of the UG, in which the focal agent is the proposer. For a game in which agent *i* is the proposer and agent *j* is the responder, *i* receives (1 – *p*_*i*_)*R*_*G*_ and *j* receives *p*_*i*_*R*_*G*_ if *p*_*i*_ ≥ *q*_*j*_, otherwise both agents receive a payoff of zero. Here, *R*_*G*_ is the resource available for game play, and is drawn from a Gaussian distribution with mean *R*_*m*_ and standard deviation σ_*R*_. In order to model limitations to an agent’s ability to accumulate resources, an agent’s total resources cannot exceed a resource cap, *R*_*C*_. Values used for *R*_*m*_ and standard deviation σ_*R*_ were based on the our previous analysis in [23] (see table S1 for the values used).

When an agent *i*’s accumulated resources, *R*_*i*_, reaches or exceeds the reproduction threshold, *R*_*T*_, it can reproduce if (i) *R*_*i*_ ≥ *R*_*T*_, (ii) the current population size, *N*, is less than the environmental carrying capacity, *K*, and (iii) there is an empty cell in the reproducing agent’s Moore neighborhood (*M* = 1). An offspring inherits the offer proportion, *p*_*i*_, and rejection threshold, *q*_*i*_, of its parent *i*. Mutations are randomly and independently introduced into an offspring *o*’s *p*_*o*_ and *q*_*o*_ at rate *r*. Mutations are randomly drawn from the uniform distributions with the range [-*m*_*r*_, *m*_*r*_]. Both the offer *p*_*o*_ and rejection threshold *q*_*o*_ are constrained to the range [0,1], and so mutations outside this range are truncated.

A parent agent invests a proportion *P* of its resources in its offspring at the time of reproduction, so that the offspring’s starting resource level *R*_*o*_ = *PR*_*i*_. While an agent can have at most one offspring per round, it may have offspring on successive rounds depending on the values for the reproductive threshold, *R*_*T*_, the amount of parental investment, *P*, the resource cap, *R*_*C*_, and its total resources*, R*_*i*_. For example if *R*_*T*_ = 100, *P* = 0.25, *R*_*C*_ = 150, and *R*_*i*_ = 145, an offspring’s initial resources would be *R*_*o*_ = 36.25, leaving the parent with *R*_*i*_ – *R*_*o*_ = 108.75, such that it space is available the agent can reproduce again on the next time step.

Each agent *i* has a fixed lifespan of *L*_*i*_ time steps, which is assigned at birth. When the agent’s age (the number of time steps since birth) reaches its lifespan, it dies and is removed from the simulation. Lifespan varies for each agent and is defined as *L*_*i*_ = *L* + *z*, where *z* is drawn from a Gaussian distribution with a mean of zero and standard deviation σ_*L*_ and then rounded to the nearest integer.

## Results

Mean offers evolved that were well above the SPNE and were consistent with data from UG experiments. The evolved mean offers were most consistent with the empirical data up to a density of *d* = 0.4 with a density *d* = 0.2 producing the overall best mean fit (see figure 1a). As population densities increased beyond 0.4, the mean evolved offers were less consistent with the empirical data (figure 1a). In contrast, in the individual-only conditions, when the effects of population structure were eliminated by randomly swapping locations of agents, offers were much lower but still far from the SPNE (figure 1a).

**Figure 1.**
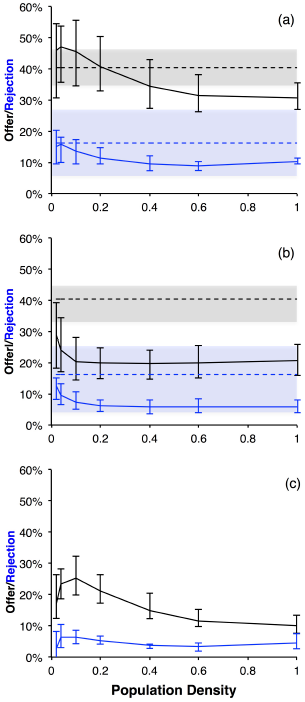
Graphs depicting the evolved mean offers (black lines) and mean rejection thresholds (blue lines) for the UG simulations averaged over all combinations of parental investment and resource caps (*P* × *R*_*C*_) plotted over population density (see figures S2, S3, S4 for individual plots for all combinations). The top figure (a) illustrates the results from the multilevel simulations, the middle figure (b) depicts the results for the individual-only simulations, and the bottom figure (c) is the difference between figures (a) and (b). The vertical interval bars depict the range of means for the 12 (*P* × *R*_*C*_) for simulation results in (a) and (b) and the vertical interval bars in (c) represent the range of differences for (a) and (b). The dark-dashed horizontal line is the empirical mean of offers from Oosterbeek and colleagues’ [2] meta-analysis of UG experiments and the gray shaded area represents the estimated standard deviation of offers. The blue-dashed horizontal line is the mean of the rejection thresholds from Oosterbeek and colleagues’ [2] meta-analysis and the blue shaded area represents the estimated standard deviation of rejection thresholds. Simulation mean offers and mean rejection thresholds that fall within the standard deviations of the meta-analysis data were considered consistent with UG experimental results.

Mean rejection thresholds evolved that were above the rejection thresholds predicted by the SPNE, but were not consistent with the evolution of Nash equilibrium strategies (figure 1a). Evolved mean rejection thresholds were the highest for the very lowest population densities (*d* = 0.02, 0.04). Unlike evolved mean offers, evolved mean rejection thresholds declined to about 11% by *d* = 0.2 and then declined to a little below 10% for higher densities. Because there is so much variation in estimated rejection threshold in the empirical data, all of the evolved mean rejection thresholds are consistent with the data, (though were biased downward) except at the lowest population densities (*d* = 0.02, 0.04).

Figure 1b illustrates the results for the individual-only conditions in which agents randomly swapped locations on each round of play. For all but the lowest densities (i.e., *d* = 0.02, 0.04) offers drop to about 20% and rejection thresholds drop to 6% on average. Although mean offer strategies do not drop to the predicted SPNE, by comparing Fig. 1a with 1b (see figure 1c) it is clear that population structure has a dramatic effect on evolved offer strategies especially for the lower population densities for *d* = 0.02, 0.1.

Figure 1c plots the differences between multilevel and individual-level simulations (see figure S4 for all individual difference plots). The greatest differences are found for population densities *d* = 0.04, 0.1. Differences for the lowest density, *d* = 0.02, are dramatically lower than for slightly higher population densities *d* = 0.04, 0.1. This effect was especially pronounced for rejection thresholds at *d* = 0.02 where for *P* = 0.75 and *R*_*C*_ = 150, the evolved mean rejection threshold for the individual-only condition was greater than the multilevel condition (see figure S4). The lowest density, *d* = 0.02, was simulated with populations of 50 agents in which small population size effects reduced the differences between multilevel and individual level simulations (figures S4).

Parental investment and resource caps influenced the evolution of mean offers and to a lesser extent mean rejection thresholds (see figure S3). Parental investment of 25% and resource cap of 100 (figure S3) resulted in the lowest mean offers. In general, parental investment of 50% or more and resource caps greater than 100 (figure S3) evolved higher mean offers.

To investigate the stability of evolved offers and how these results scale with larger populations we ran simulations with 12,500 agents (25 times larger than the 500 agent populations with *d* = 0.2) in a 250 × 250 grid for 750,000 steps. Figure 2 illustrates the dynamics of example runs for the multilevel and individual-only conditions. The mean offer strategy in the multilevel condition fluctuated over time near 40%, and the mean rejection threshold fluctuated just below 12%. Figure 3 consists of two snapshots of the same run at the start of a simulation and again at 50,000 steps by which point population structure has emerged and the population has reached a quasi-equilibrium state. Agents with low offers are represented by red. As offers increase in proportion, agents become increasingly purple and when offers strategies exceed 50% of a resource, they begin to turn blue. The range of colors in figure 3 indicates considerable variation in offer strategy. Most agents are shades of purple but there are a number of clusters of red agents and a few clusters of blue agents. These patterns gradually changed over time but the distribution of offers remained relatively stable.

**Figure 2.**
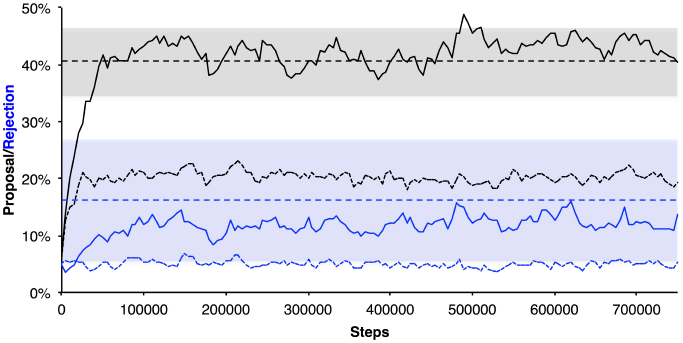
Example dynamics from a single run of 750,000 rounds with a maximum population size of *K* = 12,500, population density of *d* = 0.2, parental investment of *P* = 0.5, and resource cap of *R*_*C*_ = 150. Both multilevel and individual-level simulations are depicted using the same color scheme as figure 1. The solid lines are from the multilevel simulation and the dotted lines are from the individual-only simulation. After 50,000 rounds of play, the population in the experimental condition reached mean offers of about 40% and thereafter fluctuated close to this value. The mean rejection thresholds rose more slowly and then fluctuated close to 12%. Horizontal dashed lines represent the mean offers (black) and mean rejection thresholds (blue) from Oosterbeek and colleagues’ [2] meta-analysis.

**Figure 3.**
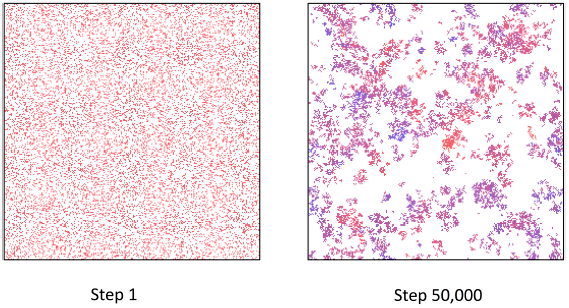
Two snapshots of a population of 12,500 agents with the same parameters as in Fig. 2. The offer strategy of each agent is represented by its color. Red agents made very low offers. As mean offers increased agents became increasingly purple. As offers increased beyond 50%, agents became increasingly blue. At step 1, agents were placed randomly in space with no population structure. As the population evolved, population structure (clusters of agents) emerged. As illustrated in the snapshot at step 50,000, clusters of agents were characterized by different offer strategies. Most clusters are shades of purple with a number of clusters of red and some clusters of blue agents.

## Discussion

Using evolutionary simulations of the UG, we have shown how population structure can facilitate the evolution of fair offers far from the SPNE, especially under conditions of low population density, while simultaneously selecting for rejection thresholds well below evolved offers, and hence contrary to assumptions of Nash equilibrium strategies. Moreover, the fair strategies that did evolve were consistent with data from UG experiments. Thus, we showed that apparently generous strategies with low rejection thresholds could robustly evolve when population structure emerges from the self-organizing behavior of agents.

Offers were the highest and most consistent with the empirical data for low to moderate population densities, in the range of d = 0.4 or less. As population density increased, evolved offers gradually decreased due to increased crowding and the decreased persistence of isolated clusters of agents resulting from crowding (figure 1a). Systematic variations in parental investment and resource storage capacity revealed that evolved offers and rejection thresholds were sensitive to these conditions as well. Evolved offers varied with population density, resource holding capacity, and parental investment, which is broadly consistent with cross-cultural findings that offers in UG experiments vary with factors such as market integration, population size, and religiosity [4]. That is, population structure as well as cultural differences in parental investment and the capacity to store resources may influence differences in levels of fairness observed across cultures (see also [29] for a recent analysis focusing on the role of network structure in the evolution of fair strategies).

Our model also predicts the considerable variation in the strategies used by participants in UG experiments. A wide range of strategies was found in simulated populations, with the precise distribution changing over time, but reflecting patterns consistent with empirical distributions. The modal offer in UG experiments is 50% with almost all other offers between 0% and 50% and a few offers above 50% [2,4]. As illustrated in figure 3, there was considerable variation in evolved offers, which changed over time but with a mean hovering near the empirical value of 40%. Most emergent clusters contained purple agents that make offers close to 50%. Fewer clusters contained red agents that offer very little and even fewer clusters contained blue agents that made offers well over 50%.

### No population structure, but still no Nash equilibrium strategies

When we eliminated the effects of population structure by swapping agent locations, mean evolved offers fell dramatically, but SPNE strategies did not evolve, nor did other Nash equilibrium strategies. In other words, offers remained substantially higher than rejection thresholds. Why should this be? One explanation is that this effect is driven by a response to variation among rejection thresholds, forcing offers to increase as a bet-hedging strategy. As we shall see, this appears to be the case, but it is not the whole story. Binmore and Samuelson [13] used a learning model of agents playing the UG with a relatively high proportion of errors in agents rejecting offers, thus simulating variation. Their results *was* a deviation from the SPNE, but was still a nearly perfect Nash equilibrium with offers around 20%.

In the individual-only version of our model, we previously showed [23] that when rejection thresholds were fixed at zero—that is, in a DG—evolved offers hovered just above zero (mutation prevents offers from staying at exactly zero). This implies that rejection thresholds force the evolution of higher offers, but the question remains as to why rejection thresholds remain so low relative to offers. In our UG model, offers and rejection thresholds were subject to random mutation. This results in a noisy evolutionary process in which the likelihood of rejected offers increases as offers and rejection thresholds converge. That is, when offers and rejection thresholds are close to each other, mutation is increasingly likely to introduce mutant rejection thresholds that are higher than offers. Because rejections are costly for both players, the convergence of offers and rejection thresholds becomes increasingly costly the closer rejection thresholds are to offers. Thus, selection acts against the convergence of offers and rejection thresholds.

High offers are selected against at the level of the individual but selection also acts against lower offers as they approach rejection thresholds. The non-Nash equilibrium strategies that evolved in our individual-only selection simulations resulted from the quasi-stabilizing forces of mutation driving up rejection thresholds, the cost of convergent offer and rejection thresholds (preventing their convergence), and individual selection against high offers. This suggests that the results from the population structure simulation experiments are a complex interplay of individual level and group level selection forces. While selection at the group level favors fairness, selection at the individual level acts both for and against fairness in the UG.

### Kin selection does not apply and the importance of resource variance

Offers far from the SPNE evolved in large part because population structure allowed spatiotemporal associations to emerge among agents with similar strategies. These associations arose because of local reproduction resulting in offspring placed adjacent to their parents, often referred to as limited dispersal. This could indicate that some type of kin selection process may have occurred. However, we believe that the classic theoretical machinery for explaining kin selection does not apply in this case. To see why, we will examine the best-case scenario for the evolution of fair strategies in the UG under conditions of population structure.

Consider two homogeneous populations of agents that play the UG for resources to reproduce, with the assumption that proposer and responder roles are always randomly assigned. One population consists of agents that always offer 50% of a resource (with *q* ≤ 0.5) and the other population consists of utterly selfish agents that offer nothing and expect nothing in return (i.e., *q* = 0). If fair agents always play selfish agents, the selfish strategy always dominates, but if fair agents only play other fair agents, then perhaps fair agents might do better than selfish agents who only play other selfish agents?

It is important to understand that this scenario differs essentially from the classic group selection argument for the evolution of altruism [30]. In the case of altruistic behavior, often modeled as a prisoner’s dilemma game, the expected fitness payoff to an individual in altruistic group is always higher than the expected payoff to an individual in a selfish group. In our scenario, in contrast, the expected payoff to individuals in either group is the same. Suppose a proposer is tasked with the division of *R*_*G*_ units of a resource. In the fair group, the proposer keeps *R*_*G*_/2 units and the responder receives *R*_*G*_/2 units; the expected payoff for members of a fair group is *R*_*G*_/2 for each game. In the selfish group, the proposer keeps *R*_*G*_ and the responder receives nothing. Because an agent is equally likely to be the proposer or the responder over time, the expected payoff is again *R*_*G*_/2. No matter the degree of generosity or selfishness of agents in homogenous groups, the expected gain in resources each interaction is always *R*_*G*_/2.

Thus, there is no difference between groups in expected resources and so apparently no fitness differences between groups. How could fairness evolve in this scenario?

This problem remains puzzling if stated in terms of Hamilton’s rule. If *p* is the proportion offered by the proposer, then a proposer offers *pR*_*G*_ and keeps (1 – *p*)*R*_*G*_. The benefit to an agent as responder is *b* = *pR*_*G*_ and the cost as proposer is *c* = *pR*_*G*_. Because selfish agents always offer less than fair agents, *p*′ < *p*, fair agents always receive less benefit when playing selfish agents and so *b* < *c*. The only hope for fair agents is to play other fair agents with the same strategy, but in that case the best that can be hope for is to break even, *b* = *c*, which violates Hamilton’s rule, *rb* – *c* > 0, even when *r* = 1 (where *r* is the probability of playing an agent with the same phenotype). Thus, Hamilton’s rule is of no help in explaining the evolution of fairness. Nevertheless, fair offers robustly evolved, which implies that some group property is missing from the analysis.

Returning to our two groups of agents, there is one factor in which the groups differ: the *variance* in resources among individuals within a group. In fair groups, proposers offer an even split. If we assume on each round of play that proposer and responder roles are randomly assigned and that proposers find exactly *R*_*G*_ units of a resource, then as we saw above the expected payoff for fair or selfish players in any homogenous group is *R*_*G*_/2. After one round of play, however, although there is zero variance in the resources accumulated by agents in fair groups, there is considerable variance in selfish groups as given by the following equation.

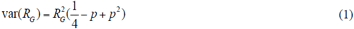

If agents are fair (*p* = 0.5) and half the agents in a population are proposers and the other half responders, the variance in resources after the members of the group play a round is zero. As *p* approaches zero (selfish) or one (selfless), the variance increases rapidly. The magnitude of variance in resources is the only difference between fair and selfish groups, and so must provide an explanation for the evolution of fairness in the context of our model.

Differences between groups in resource variance create the opportunity for differences in the efficiency of the conversion of resources into offspring, at the level of the group, when there are constraints on this flow. In our model, the benefit to fair offers comes from the more efficient flow of resources into the production of offspring. In contrast, greater variance among individuals in selfish groups creates inefficiencies in the conversion of resources into offspring. We introduced two important assumptions that constrained the conversion of resources into offspring: population carrying capacity and limited spatial availability. Agents could only produce offspring if the total number of agents, *N*, was less than the maximum population size *K* (*N* < *K*) and if an empty nearby cell was available. Since these assumptions resulted in agents sometimes waiting to reproduce, lucky selfish agents accumulated abundant resources that they could not fully convert to offspring—resources that in fair groups were partially distributed to other neighbors (see figure 4). Thus, because fair agents more evenly distributed resources, agents in fair groups more efficiently converted resources into offspring as illustrated in figure 4. This implies that an analysis of the fitness costs and benefits of fairness for individuals requires an analysis of the flow of resources at the level of population structure and not merely an accounting of benefits at the individual level.

**Figure 4.**
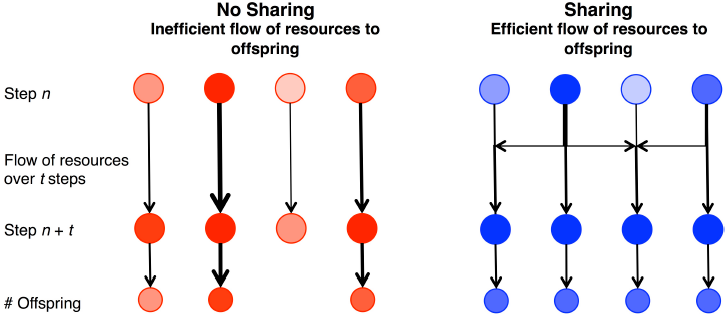
A simplified illustration of the flow and conversion of resources over a sequence of games resulting in the differential production of offspring as a function of strategy. Fair agents are blue (*i.e., p* = 0.5, *q* ≤ 0.5) and selfish agents (*p* = 0, *q* = 0) are red. Shading represents resource status. For selfish agents (left), reproduction of offspring depends only on their chance accumulation of resources over *t* steps. For fair agents (right), reproduction depends on both their chance accumulation of resources and the sharing of resources among each other.

### Biological implications

Similar constraints are universal in biological populations. For example, a fundamental constraint on the conversion of resources into offspring is imposed by development. In mammals, gestation periods vary across species, but gestation and parental care impose constraints on how rapidly individuals reproduce offspring. This is a developmental constraint on how fast animals can reproduce [31]. For example, vampire bats give birth to a single offspring about once a year [32]. A vampire bat cannot increase its rate of birth to twice a year by consuming twice as much blood. Feeding a female vampire twice as much blood as she can use and store will not double her reproductive rate, but rather is a waste of blood from the perspective of fitness, and indeed may be part of why many organisms exhibit Type II functional responses to resources [33]. However, if the extra blood is shared with others, that excess blood can be transformed into offspring by increasing the survival of starving bats (Wilkerson, 1984) [34]. Indeed, vampire bats share blood meals within groups even with non-kin [34,35].

The aim of our modeling was to not only explain fair offers in the UG far from SPNE but also to explain the low rejection thresholds characteristic of experimental data from UG experiments. We found that generous offers with low rejection thresholds evolved in structured populations and that factors such as parental investment and the capacity to store resources affected the degree of fairness that evolved. No single globally-adopted strategy evolved in any of our simulations. Instead, populations were composed of a distribution of strategies that on average fluctuated around mean values consistent with those seen empirically. This population variation may be inherent in many evolutionary processes—biological or cultural—and may explain the within-population variation typical of UG experiments. Finally, analysis of the flow of resources through structured populations may shed light into the evolution of cooperation more generally as well as the evolution of other types of social behavior and group-level organization.

## Methods

In simulations of our model as described above, limited dispersal and local mobility produced spatial assortment by phenotype. Because group structure is known to affect selection on social traits [35-41], we term these “multilevel simulations.” In order to assess the effects of assortative group structure, we additionally paired each multilevel simulation run with a control simulation in which each agent swapped location with another randomly chosen agent on each round of play. Following [23], we term these “individual-only simulations,” as they allowed us to compare the effects of population structure with unstructured random interactions among agents while preserving spatial-group structure. In other words, selection is at the individual level only, but we maintain spatial-group structure so that interactions do not vary between conditions. Most simulations were run for 100,000 time steps, which was sufficient time for the evolved offers and rejection thresholds to reach long-term values (we also ran individual simulations with much larger populations and for 750,000 steps to investigate scalability and stability; see below).

Initial population size *N* was always set to its maximum size *K.* Each agent, *i*, was randomly placed at a unique random location in the 2D grid space. Initial offers, *p*_*i*_, and rejection thresholds, *q*_*i*_, were each randomly drawn from a uniform distribution [0, 0.1], resulting in very selfish agents with fairly low rejection thresholds at the beginning of all simulations. All other initial parameter values are described in table S1.

We systematically varied three parameters: population density, *d* (the size of the grid divided by *K*), parental investment, *P*, and the resource cap, *R*_*C*_. Less crowded population structures allow greater isolation of groups, which facilitates stronger selection at the group level. A total of 100 simulations were run for each set of parameter values (see Table S1).

## Acknowledgments

We would like to thank the Institute for Advanced Studies in Glasgow, Scotland for the series of workshops on “Limits to Rationality in Financial Markets” (and the organizers: Michael Grinfeld, Harbir Lamba, and Rod Cross) during which the first versions of the models reported here were developed. We also thank Bert Baumgaertner, Linnda Caporael, Sabine Durand, Michael Grinfeld, Jay Jefferson, Thomas Kohnle, and Elizabeth Matthews for their comments on versions of this draft. Matt L. Miller was supported in part by the National Science Foundation Graduate Research Fellowship Program under Grant no. 1650042.

## Supplementary Information

**Table S1.** Fixed parameters, initial conditions, and parameter sweeps.

| Description | Parameter | Values and/or descriptions |
| --- | --- | --- |
| **Fixed Parameters** |  |  |
| Game Resources | $R_m$ | 10 – the average game resource played for each step |
| Game Resources Standard Deviation | $\sigma_R$ | 1 – the standard deviation for $R_G$ |
| Expected Life Span | $L$ | 100 – time steps |
| Life Span Standard Deviation | $\sigma_L$ | 5 – the standard deviation for $L$ |
| Reproduction Resource Threshold | $R_T$ | 100 – the resource level required for an agent to reproduce. |
| Mutation rate | $r$ | .01 |
| Mutation range | $m_r$ | 0.25 – e.g., if $p = 0.1$, then minimum and maximum values possible are 0 and 0.35). Mutations were uniform random within the range $[-m_r, m_r]$. |
| Grid Size | $X \times Y$ | $50 \times 50$ – spatial dimensions in cells |
| Moore Radius | $M$ | 1, 2 – depending on context (UG, reproduction, search) |
| **Initial conditions** |  |  |
| Starting Resource | $R_0$ | 50 – the amount of resources an agent starts with at the beginning of a simulation (varies for agents born later in a simulation by parental resources and investment), |
| Age | $L_0$ | 1 – age of agents at the start of a simulation |
| Population size | $N_0$ | set to the maximum population size |
| Agent $i$: | | |
| offer proportion | $p_i$ | $[0, .1]$ – drawn from uniform-random distribution |
| rejection threshold | $q_i$ | $[0, .1]$ – drawn from uniform-random distribution |
| **Parameter Sweeps** |  |  |
| Max Population Size | $K$ | 50, 100, 250, 500, 1000, 1500, 2500 |
| Parental Investment | $P$ | .25, .5, .75 |
| Resource Cap | $R_C$ | 100, 150, 300, unlimited |

**Figure S1.**
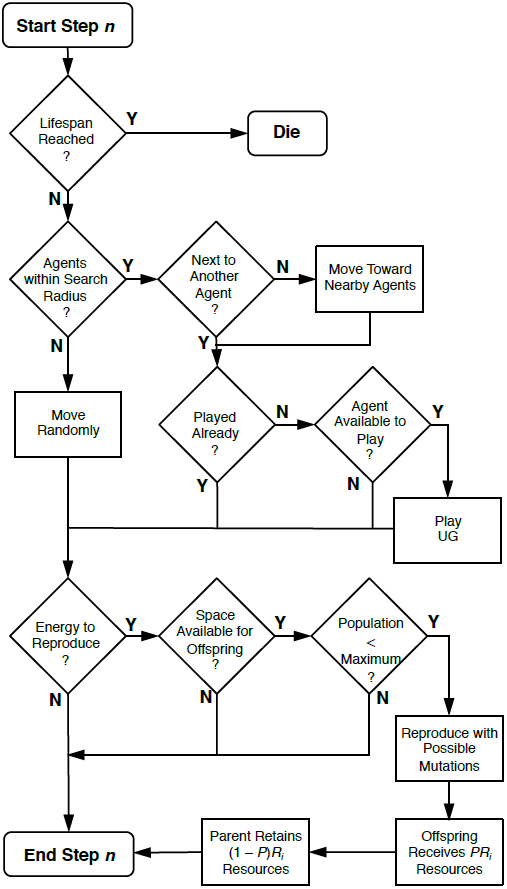
A flow chart of an agent *i*’s decisions and key events on each round of play. If an agent does not die, it begins by searching for at least one other agent to play with. If there is no other agent in its Moore search radius (*M* = 2), it moves in a random zigzag pattern on each step till it contacts a least one other agent. When it is next to another agent and both agents have not yet played, they play the UG. Whether or not an agent played, it can attempt to reproduce if it has sufficient resources, there is an open cell in its Moore neighborhood (*M* = 1), and the population size is less than *K*. Offspring are placed in a randomly selected vacant cell in the agent’s Moore neighborhood (*M* = 1). If the agent reproduces, then mutations for *p*_*i*_ and *q*_*i*_ each occur with probability *r* resulting in the offspring’s *p*_*o*_ and *q*_*o*_. The offspring agent receives *R*_*o*_ = *PR*_*i*_ resources from the parent and the parent retains *R*_*i*_ – *R*_*o*_ resources (see text and table S1 for more detail).

**Figure S2.**
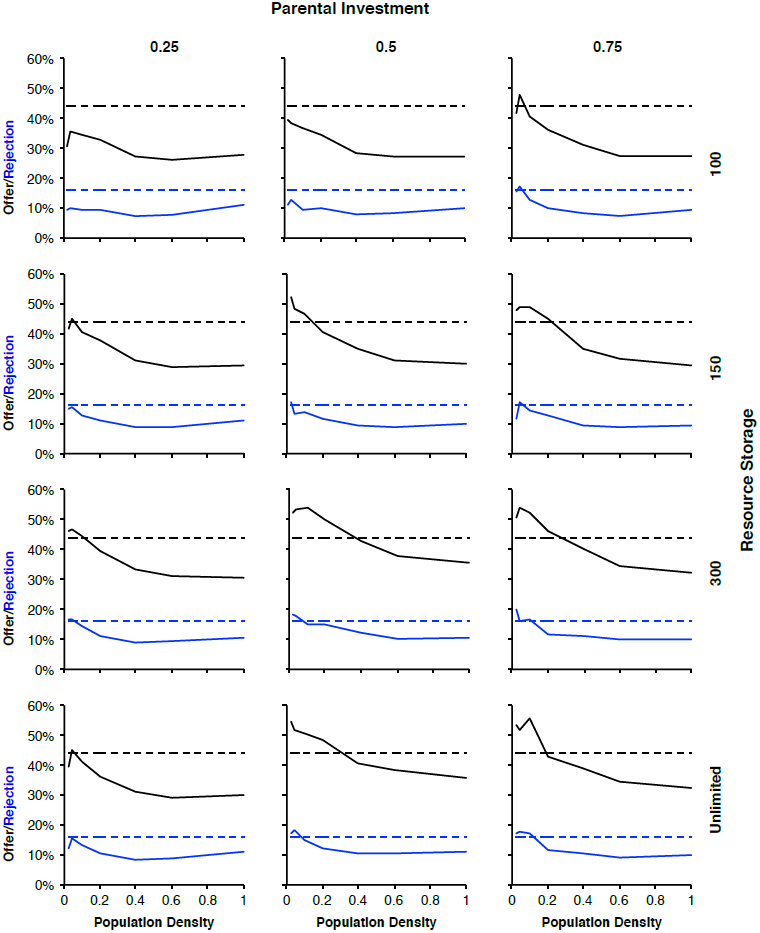
The effects of parental investment and resource storage capacity on evolved offer means and rejection threshold means as a function of population density for all simulation conditions. Columns illustrate parental investment levels of 0.25, 0.5, and 0.75 and rows depict resource storage capacities of 100, 150, 300, and unlimited. Solid black lines are the evolved offer means and solid blue lines are the evolved rejection thresholds. Dashed lines are the corresponding empirical means for offers (black) and rejection thresholds (blue; see figure 1 in text for the empirical standard deviations). As population density increases, group population structure through crowding, begins to break down and offers tend to decrease.

**Figure S3.**
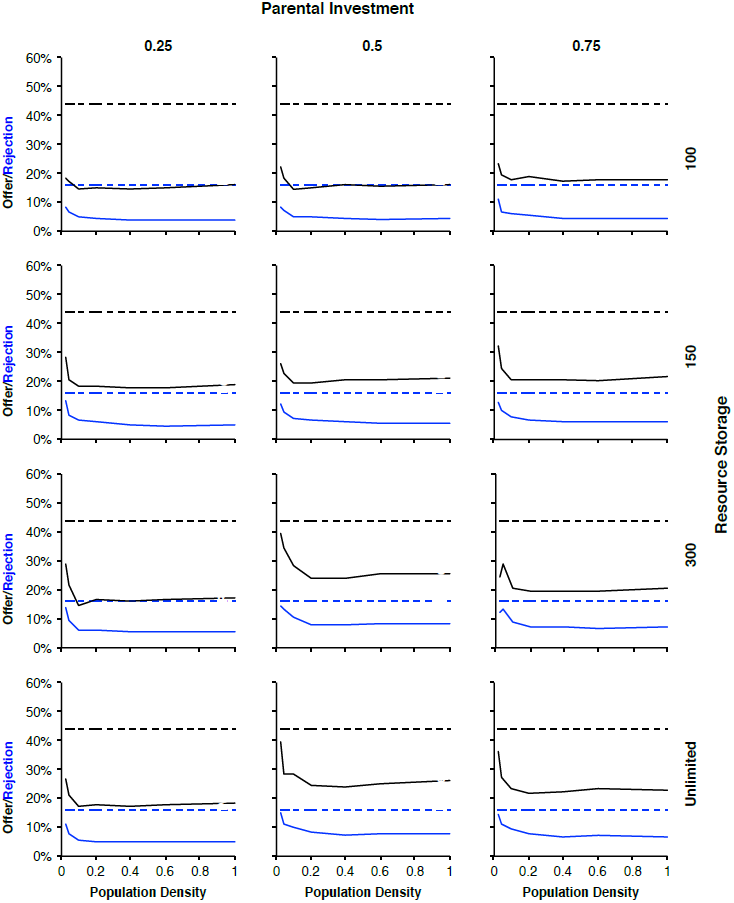
The effects of parental investment and resource storage capacity on evolved offer means and rejection threshold means as a function of population density for all simulation conditions when agents randomly swapped locations on each step. As in figure S3, columns illustrate parental investment levels of 0.25, 0.5, and 0.75 and rows depict resource storage capacities of 100, 150, 300, and unlimited. Solid black lines are the evolved offer means and solid blue lines are the evolved rejection thresholds. Dashed lines are the corresponding empirical means for offers (black) and rejection thresholds (blue; see figure 1 in text for the empirical standard deviations). Both mean offers and rejection thresholds are highest for the lowest population density simulated (0.02; i.e., 50 agents). Small population effects can lead to populations drifting to higher levels, which largely explains the higher evolved values that rapidly drop off with increasing population size.

**Figure S4.**
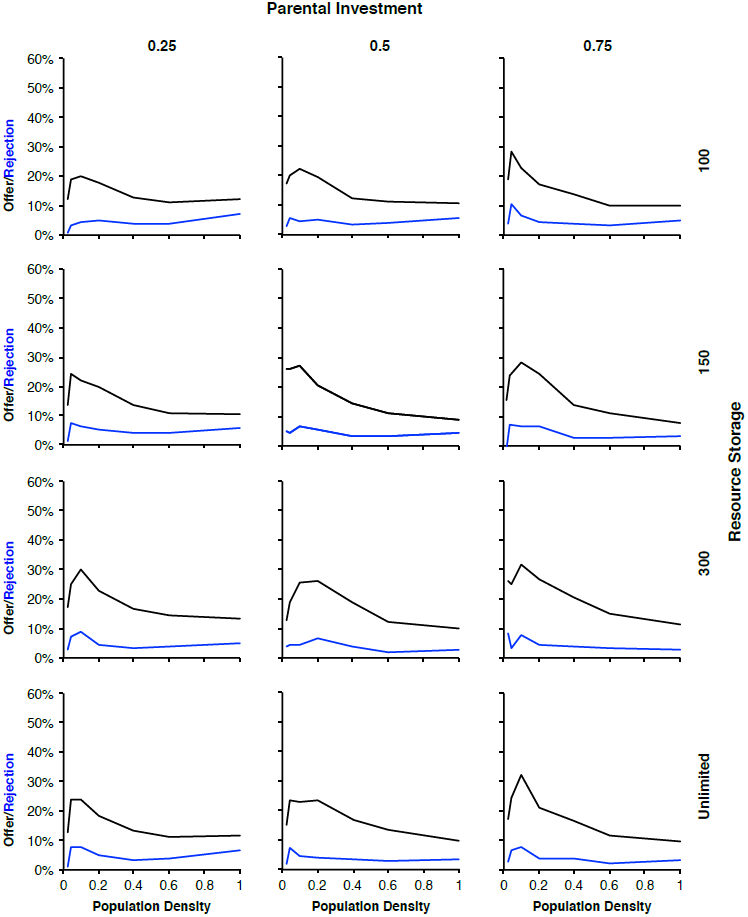
Differences between multilevel simulations (figure S2) and individual level-only simulations (figure S3). As with figures S2 and S3, columns illustrate parental investment levels of 0.25, 0.5, and 0.75 and rows depict resource storage capacities of 100, 150, 300, and unlimited. Solid black lines are the evolved offer means and solid blue lines are the evolved rejection thresholds. For most conditions, the greatest difference occurs for population densities of 0.04 (100 agents) and 0.1 (250 agents). At a population density of 0.02 (50 agents), small population effects are important resulting in smaller differences. As population density increases from 0.1, the differences between mean offers (figure S2 – figure S3) tend to decrease. However, the differences between rejection thresholds (figure S2 – figure S3) exhibit non-monotonic behavior initially decreasing and often increasing slightly at higher densities.

